# Sprouty2 limits intestinal tuft and goblet cell numbers through GSK3β-mediated restriction of epithelial IL-33

**DOI:** 10.1101/2020.03.09.984369

**Authors:** Michael A. Schumacher, Jonathan J. Hsieh, Cambrian Y. Liu, Keren L. Appel, Amanda Waddell, Dana Almohazey, Kay Katada, Jessica K. Bernard, Edie B. Bucar, Safina Gadeock, Kathryn M. Maselli, M. Kay Washington, Tracy C. Grikscheit, David Warburton, Michael J. Rosen, Mark R. Frey

**Affiliations:** The Saban Research Institute, Children’s Hospital Los Angeles, Los Angeles, CA 90027; Department of Pediatrics and Department of Biochemistry and Molecular Biology, University of Southern California Keck School of Medicine. Los Angeles, CA 90089; Division of Gastroenterology, Hepatology and Nutrition, Cincinnati Children’s Hospital Medical Center, Cincinnati, OH 45229; Department of Stem Cell Research, Institute for Research and Medical Consultation (IRMC), Imam Abdulrahman Bin Faisal University, Dammam, Saudi Arabia; Departments of Pathology, Microbiology, and Immunology, Vanderbilt Ingram Cancer Center, Vanderbilt University Medical Center, Nashville, TN 37232; Department of Surgery, Keck School of Medicine, University of Southern California, Los Angeles, CA, 90089; Department of Pediatrics, University of Cincinnati College of Medicine, Cincinnati, OH 45229

**Keywords:** Sprouty2, IL-33, IL-13, tuft cell, goblet cell, colitis

## Abstract

Dynamic regulation of intestinal cell differentiation is crucial for both homeostasis and the response to injury or inflammation. Sprouty2, an intracellular signaling regulator, controls pathways including PI3K and MAPKs that are implicated in differentiation and are dysregulated in inflammatory bowel disease. Here, we asked whether Sprouty2 controls secretory cell differentiation and the response to colitis. We report that colonic epithelial Sprouty2 deletion led to expanded goblet and tuft cell populations. Sprouty2 loss induced PI3K/Akt signaling, leading to GSK3β inhibition and epithelial interleukin (IL)-33 expression. In vivo, this resulted in increased stromal IL-13+ cells. IL-13 in turn induced tuft and goblet cell expansion in vitro and in vivo. Sprouty2 was downregulated by inflammation; this appeared to be a protective response, as VillinCre;Sprouty2^F/F^ mice were resistant to DSS colitis. In contrast, Sprouty2 was elevated in colons of inflammatory bowel disease patients, suggesting that this protective epithelial-stromal signaling mechanism is lost in disease.

## Introduction

Tight control of secretory cell differentiation is crucial for maintaining a healthy intestinal epithelium. For example, secretory tuft and goblet cells perform defensive roles against disease by promoting beneficial “weep and sweep” type immune responses and providing a physical mucus barrier against the external environment^1–3^, and defects in these cells predispose to or exacerbate barrier dysfunction and inflammation^4,5^. On the other hand, excessive secretory cell development at the expense of absorptive lineages could in theory also be detrimental. Like many elements of intestinal physiology, a careful balance in a “Goldilocks zone” is required. Furthermore, the optimal mix of lineages may change rapidly in the face of injury or inflammation^6–8^, requiring the organ to rapidly alter its cellular composition to deal with current conditions.

Despite the importance of tight, and apparently dynamic, regulation of secretory differentiation, relatively little is known about how this balance is maintained. What is clear is that intracellular signaling plays a major role. PI3K and MAPK signals rapidly alter the balance of secretory cell development^9–11^. Some of these pathways are dysregulated in inflammatory bowel disease (IBD), and defects in them likely play a key role in epithelial dysfunction during chronic inflammation^12,13^. However, the mechanisms by which diverse signals (MAPKs, PI3K/Akt, Wnts, etc.) are integrated and regulated in the gut are not well-understood.

Sprouty2 is a widely-expressed regulator and integrator of intracellular signaling downstream of receptor tyrosine kinases. It was originally identified as a critical player in branching morphogenesis in the developing mammalian lung^14–17^, and its ability to bind and inhibit Raf^18,19^ placed it as a regulator of the canonical MAPK cascade (i.e. Ras-Raf-MEK-ERK). Since then, docking sites on Sprouty2 for molecules such as the ubiquitin ligase Cbl^20,21^, the adapter protein Grb2^22^, Src^22,23^, and components of the PI3K pathway^24,25^ have been identified, pointing to a broader and more integrative role in controlling and shaping signaling. As these pathways are involved in cellular differentiation and immune responses in various tissue types, we sought to determine the function of Sprouty2 in the colonic epithelium.

To address this question, we used coordinated in vivo (a novel intestinal epithelial-specific Sprouty2 knockout mouse) and in vitro (colonoid culture) approaches to study how Sprouty2 fine-tunes the intestinal environment by modulating epithelial function, cell differentiation, and cytokine production. We find that Sprouty2 is a key regulator of tuft and goblet cell census in the colon, through an epithelial-stromal circuit involving PI3K, GSK3β, IL-33, and IL-13. Furthermore, Sprouty2 was highly responsive to acute inflammatory stimuli, suggesting that it is a mediator of dynamic changes to secretory cell populations in response to challenge. While inflammatory cytokine-induced suppression of Sprouty2 was conserved in healthy human tissues, Sprouty2 levels were in contrast elevated in human IBD. Thus, loss of this mechanism potentially contributes to disease by removing a compensatory response to injury and inflammation.

## Results

### Sprouty2 is robustly expressed in the colonic epithelium but is repressed by inflammation

As a first step toward identifying the function of Sprouty2 in the gut, we analyzed its expression pattern by qPCR, in situ hybridization, and western blot. Sprouty2 was expressed throughout the intestinal epithelium, with highest levels in the colon (Figure 1a). Levels were low in the sub-epithelial space (Figure 1b). As an integrator of extracellular stimuli, Sprouty2 expression is repressed by inflammatory challenge in airway epithelial cells^26^; to test if it is similarly regulated in the colon, we subjected mice to acute DSS colitis (Figure 1c). Sprouty2 levels were significantly reduced by 3 days post-DSS injury (Figure 1d), correlating with elevated TNF levels (Figure 1d). In vitro, TNF inhibited Sprouty2 expression in murine colonic epithelial organoid cultures (colonoids), and in cultured mouse (YAMC) colonocytes (Figure 1e and f). Collectively, these data show that Sprouty2 is an inflammation-regulated epithelial target both in vivo and in vitro.

**Figure 1.**
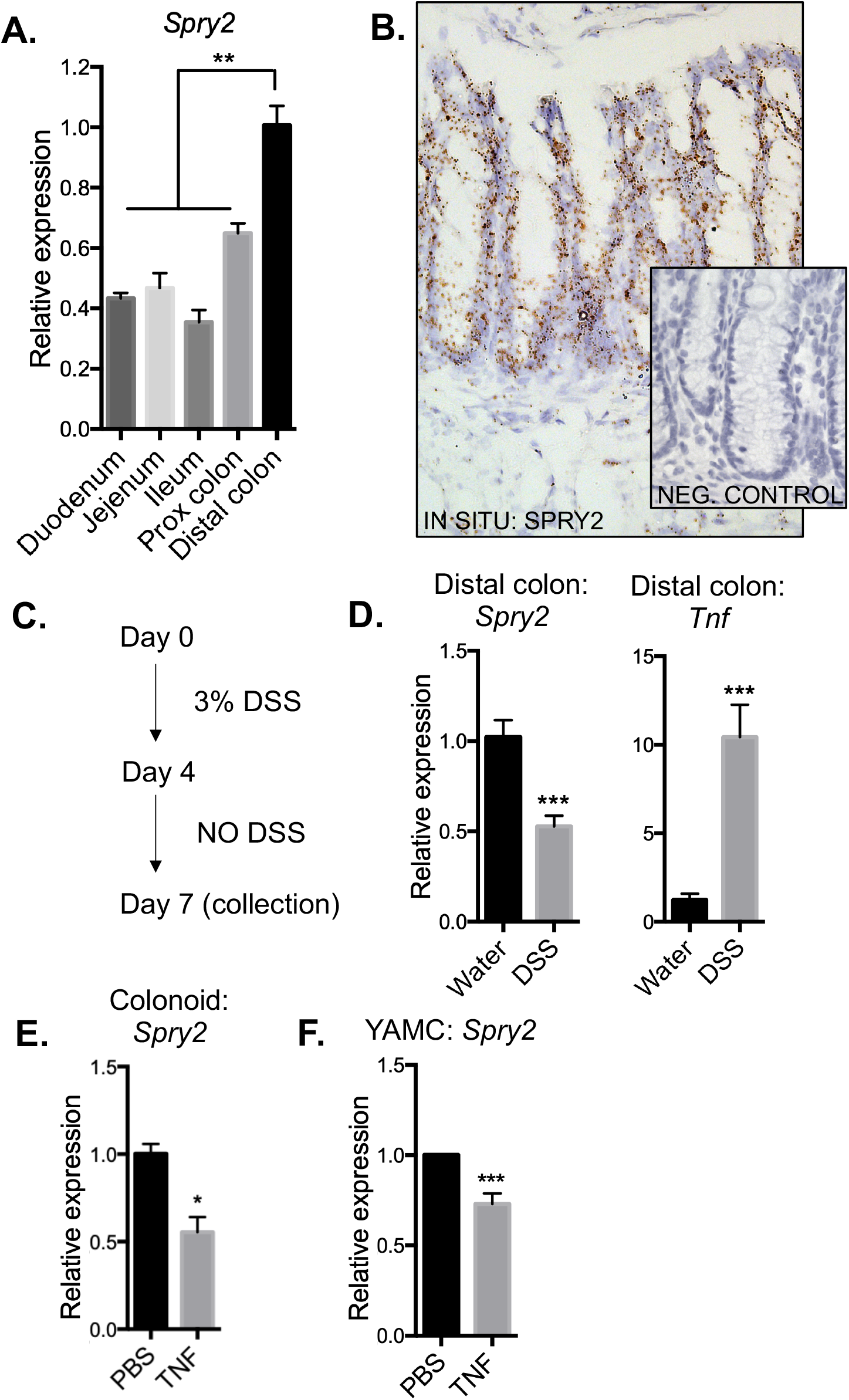
Sprouty2 is highly expressed in the colonic epithelium and repressed by inflammation. (A) Expression of *Sprouty2* was analyzed along the length of the intestinal tract by qPCR. (B) In situ analysis of murine distal colonic sections showed *Sprouty2* is predominantly expressed within the epithelial compartment. (C) Mice were subjected to 3% DSS in drinking water for 4 days followed by 3 days chase to induce colitis and (D) analyzed by qPCR for *Sprouty2* and *Tnf*; n=9-10 mice per group. (E) Primary murine colonic epithelial colonoid cultures and (F) immortalized mouse colon cells (YAMC) were treated with TNF (100 ng/ml) for 24 hours and analyzed by qPCR for *Spry2* levels. n=3-6 independent experiments. *, *p*<0.05; **, *p*<0.01; ***, *p*<0.001.

### Tuft and goblet cell numbers are increased in the colons of Spry2^IEKO^ mice

To understand the functional outcome of inflammation-induced Sprouty2 repression in the colonic epithelium, we generated intestinal epithelial-specific Sprouty2 deletion mice (VillinCre;Sprouty2^flox/flox^, hereafter termed Spry2^IEKO^; Figure 2a). Since the MAPK and PI3K cascades regulated by Sprouty2 are intimately involved in differentiation^9,11^ we analyzed lineage markers (Figure 2b-f) and found a significant increase in tuft (*Dclk1*, *Trpm5, Il25*) and goblet (*Muc2* and *Tff3*) cell marker expression in colonic tissue of Spry2^IEKO^ mice compared to Cre-negative littermate controls. In contrast, markers for enteroendocrine (*ChgA*), stem (*Lgr5*, *Lrig1*), and absorptive (*Car2, Aqp8)* cells did not change, suggesting specificity in the response. To confirm that RNA changes correlated with increased tuft and goblet cell numbers, we stained colonic sections for Dclk1, Muc2, and ChgA (Figure 2g-i). Spry2^IEKO^ mice showed an increase in the number of both tuft and goblet cells, but not enteroendocrine cells. The differential regulation of secretory cells suggests an additional modulator of secretory pathways may be required for enteroendocrine specification^27^. As secretory cells have been linked to barrier integrity maintenance^28^, we also tested whether Spry2 deletion modulated epithelial barrier function. Spry2^IEKO^ mice exhibited reduced barrier permeability (Figure 2j), demonstrating a functional change driven by loss of Sprouty2.

**Figure 2.**
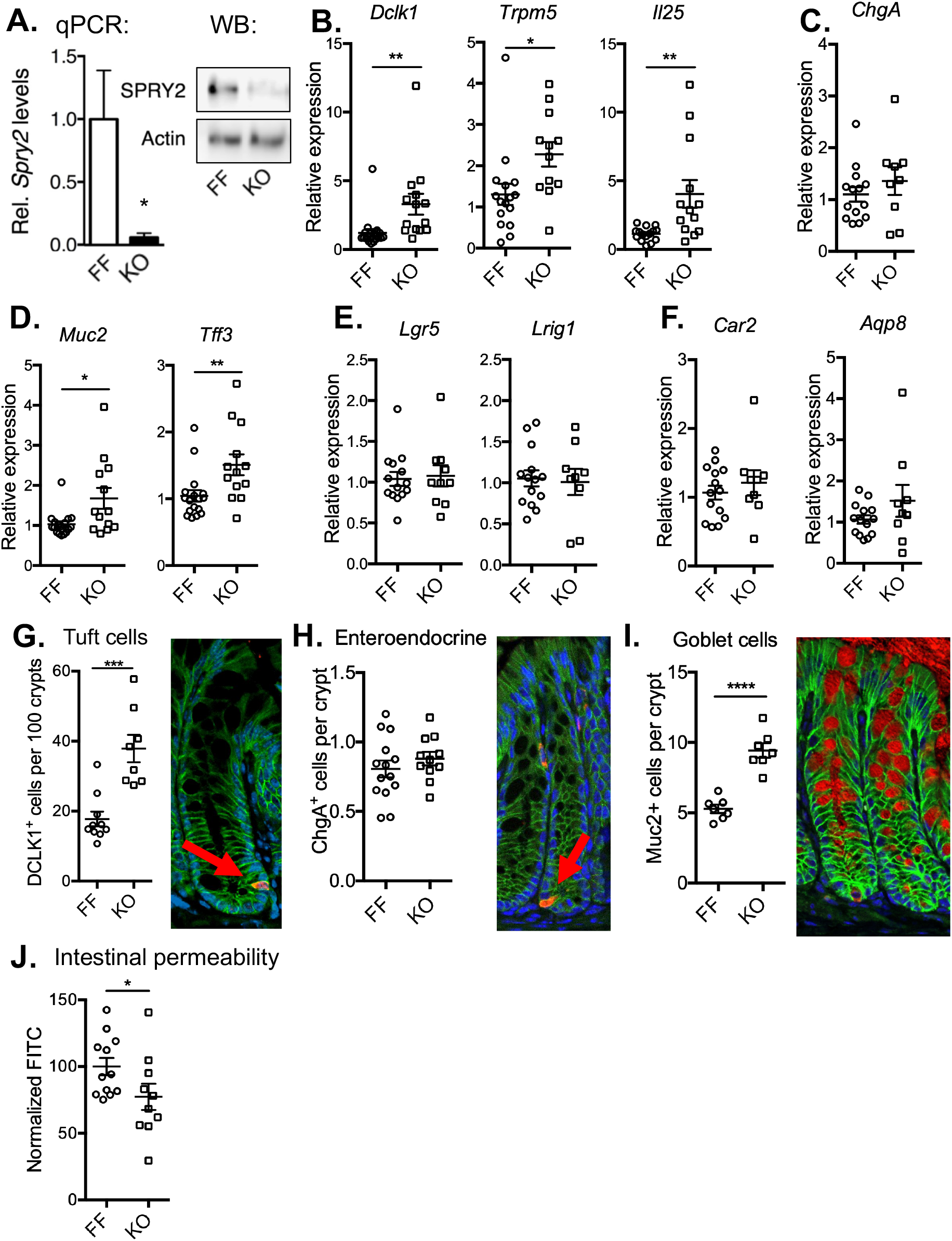
Loss of epithelial Sprouty2 results in expansion of colonic tuft and goblet cell numbers. (A) Deletion of Sprouty2 in Spry2^IEKO^ mice was confirmed by qPCR and western blot of colonic homogenates. (B) Tuft (*Dclk1*, *Trpm5*, *IL-25*), (C) enteroendocrine (*Chga*), (D) goblet (*Muc2*, *Tff3*), (E) stem (*Lgr5*, *Lrig1*), and (F) absorptive enterocyte (*Car2*, *Aqp8*) markers were measured in Spry2^IEKO^ colons by qPCR; n=10-14 mice per group. (G) Tuft (Dclk1^+^), (H) enteroendocrine (ChgA^+^), and (I) goblet (Muc2^+^) cells were quantified by immunofluorescence staining of distal colonic sections; n=7-14 mice per group. (J) Intestinal permeability was quantified by gavage of 4kDa-FITC and blood collection after 4 hours; n=10-12 mice per group. *, *p*<0.05; **, *p*<0.01; ***, *p*<0.001; ****p<0.0001.

### Sprouty2 deletion induces colonic epithelial IL-33 expression

The intestinal epithelium is a potent producer of cytokines that shape immune responses in the gut. IL-33 is a member of the IL-1 super-family with reported expression in both the epithelium and mesenchyme. It is linked to secretory cell development, which may be a response mechanism to protect the epithelium from insult. However, the environmental and intracellular signals differentially regulating IL-33 expression in different cell types are not well understood. To test whether Sprouty2 might influence tuft and goblet cell expression through modulation of IL-33 expression, we first performed qPCR analysis on colonic mucosa. We observed a significant increase in *Il33* expression in Spry2^IEKO^ mice versus control littermates (Figure 3a). No changes in baseline *Tnf* or *Il10* were observed (Figure 3a) and other immune cytokines were undetectable (*Cxcl2*, *Il4*, not shown). We confirmed IL-33 induction at the protein level by ELISA on epithelial isolates (Figure 3b). Similar to in vivo results, *Il33* was durably elevated in passaged knockout colonoids (Figure 3c), specifically demonstrating increased epithelial expression. Interestingly, while tuft and goblet cell markers were elevated in Sprouty2-null epithelia in vivo (Figure 2), this was not the case in passaged colonoids (Figure 3c), suggesting a requirement for an additional, extra-epithelial signal.

**Figure 3.**
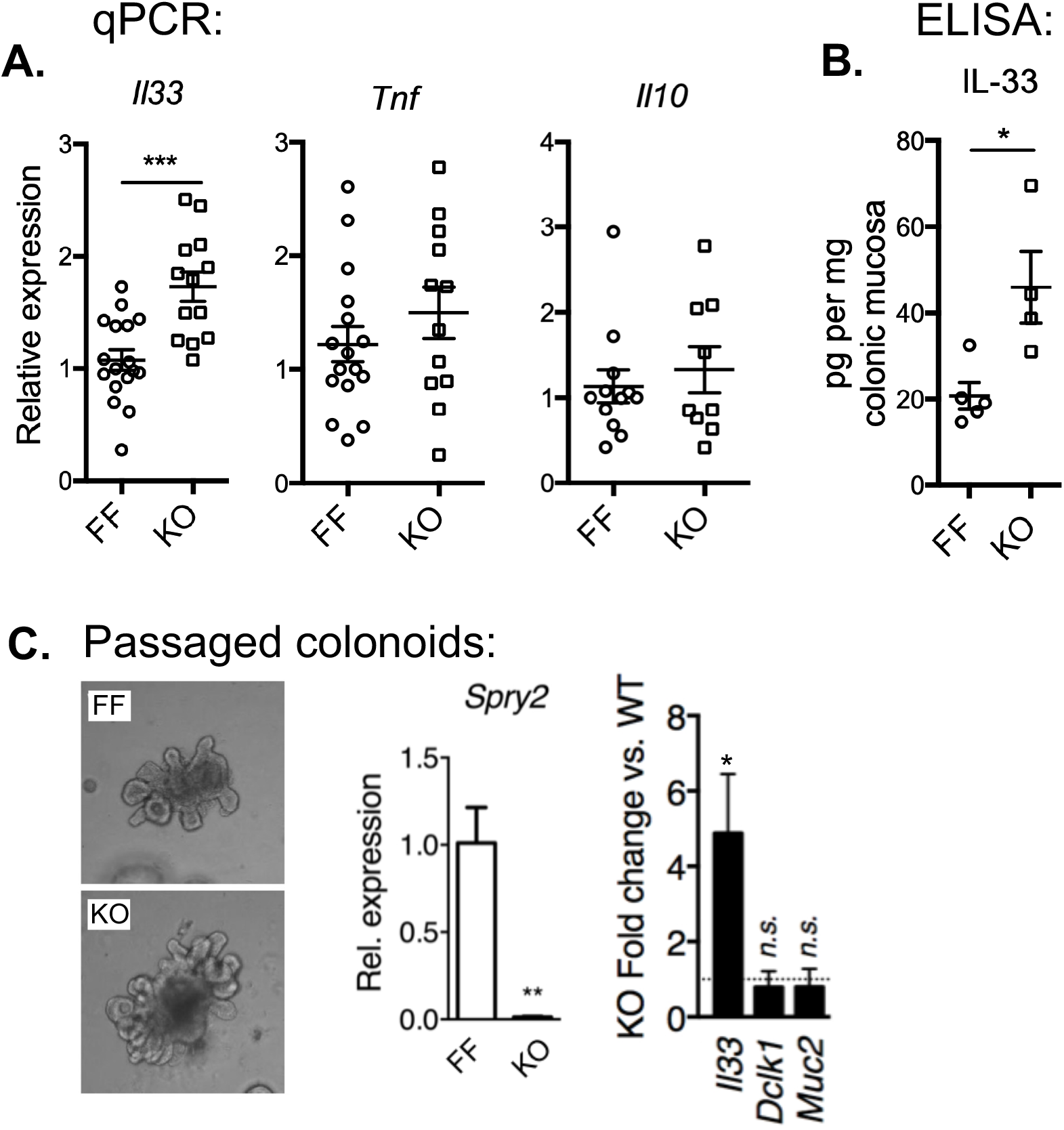
Colonic epithelial IL-33 expression is increased in Spry2^IEKO^ mice. (A) *Il33*, *Tnf*, *and Il10* levels were measured by qPCR in colonic homogenates from Spry2^FF^ and Spry2^IEKO^ mice; n=9-16 mice per group. (B) IL-33 protein was measured in colonic mucosa by ELISA; n=4-5 mice per group (C) Expression of *Spry2*, *Il33*, *Dclk1*, *and Muc2* was assayed by qPCR in passaged colonoids generated from Spry2^FF^ and Spry2^IEKO^ mice. n=3-6 independent experiments. *, *p*<0.05; ** *p*<0.01; ***, *p*<0.001.

### Tuft and goblet cell expansion in Spry2^IEKO^ mice requires stromal signals

In the intestine^1,6,29^, stomach^30^ and skin^31^ IL-33 can drive IL-13 release from type-2 innate lymphoid cells (ILC2s). We performed in-situ analysis for IL-13^+^ cells and found a significant increase in their numbers within colonic stromal tissue of Spry2^IEKO^ mice (Figure 4a). Similarly, we found elevated levels of *Alox5*, a marker expressed by ILC2s^32,33^ (Figure 4b).

**Figure 4.**
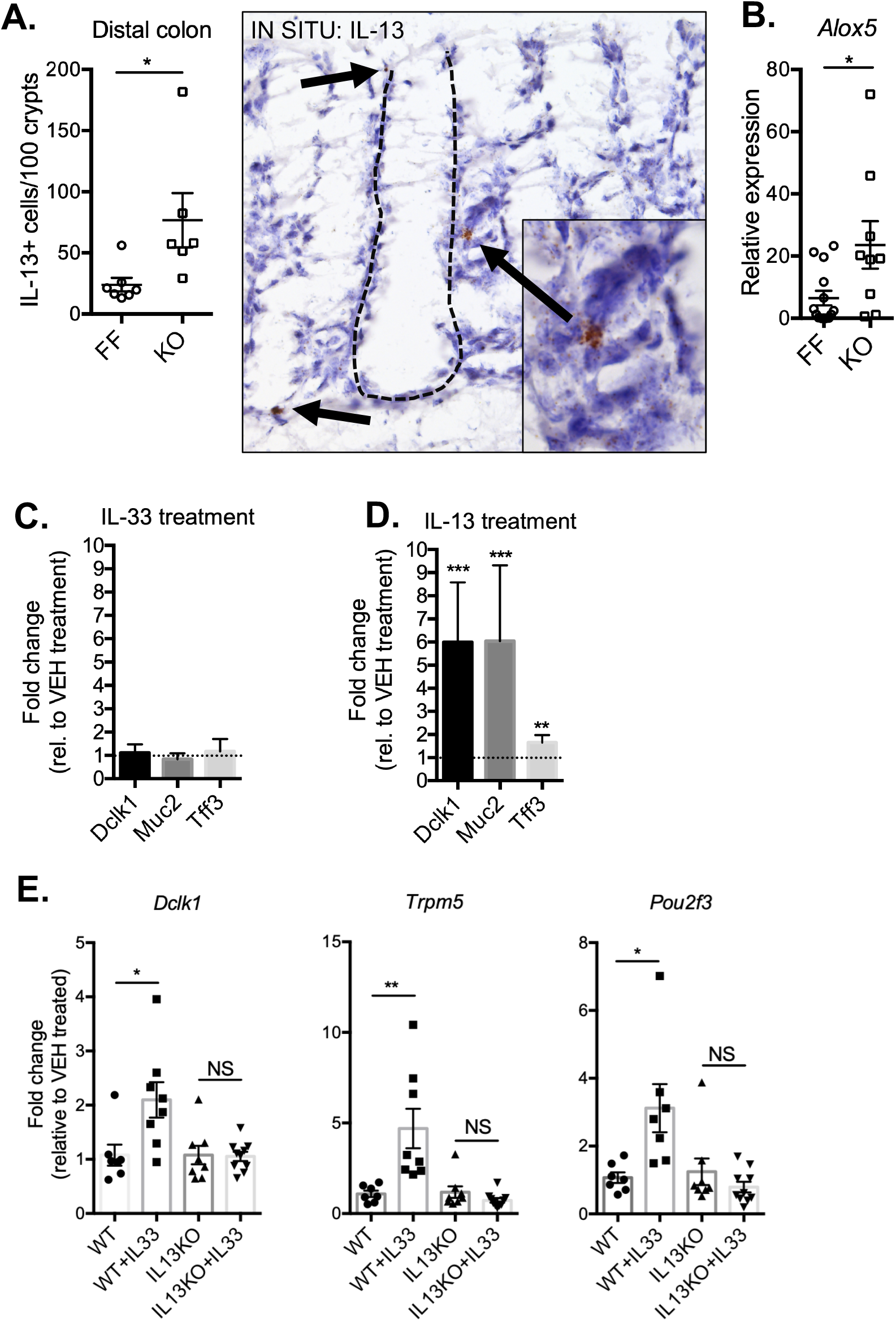
IL-33-driven tuft and goblet cell expansion requires stromal IL-13. (A) Distal colonic sections from Spry2^FF^ and Spry2^IEKO^ mice were probed for *Il13* by in situ RNAScope analysis and counted for IL-13^+^ cells per 100 crypts (dashed line, base of epithelium); n=6-7 mice per group. (B) Expression of ILC2-signature gene, *Alox5*, was quantified by qPCR of distal colonic tissue. Colonoids generated from WT mice were treated with (C) IL-33 (100 ng/ml) or (D) IL-13 (10 ng/ml) for 24 hours, and analyzed for tuft (*Dclk1*) and goblet cell (*Muc2, Tff3*) marker expression; n=5 independent experiments. (E) WT and IL-13^-/-^mice were i.p. injected with PBS or IL-33 for 4 days, and distal colonic expression of tuft cell markers (*Dclk1, Trpm5, Pou2f3*) was assayed, revealing a requirement for IL-13 in IL-33 mediated tuft cell induction in the colon; n=7-10 mice per group. *, *p*<0.05; **, *p*<0.01; ***, *p*<0.001.

To test if tuft and goblet cell expansion is an epithelial-intrinsic response to IL-33 or in contrast requires IL-13, we treated colonoids from wild-type mice with these cytokines. Exogenous IL-33 was not sufficient to promote expression of tuft *(Dclk1)* or goblet *(Muc2, Tff3)* cell markers in colonoids (Figure 4c), while IL-13 robustly induced these genes (Figure 4d). In vivo, we found that tuft cell marker induction in the colon by IL-33 injection did not occur in IL-13KO mice (Figure 4e), extending previous work demonstrating similar findings for goblet cells^34^. Together these results suggest an epithelial-stromal circuit in which Sprouty2 downregulation allows for IL-33 expression in the epithelium, promoting IL-13 release in the sub-epithelial cells which in turn signals back to the epithelium to alter cell differentiation patterns, consistent with a recent study on IL-33 driven ileal goblet cell hyperplasia^34^.

### GSK3β inhibition is driven by Sprouty2 loss, leading to IL-33 expression

To understand the precise signaling mechanisms through which Sprouty2 controls epithelial IL-33, we performed a screen for phosphorylated growth factor receptor signaling outputs on Spry2^IEKO^ and Cre-negative Spry2^flox/flox^ littermates (R&D Systems, Proteome Profiler; not shown). This analysis showed increased Serine 9 phosphorylation on the Wnt regulator, GSK3β, in Spry2^IEKO^ mucosa. S9 is an Akt-mediated inhibitory phosphosite on GSK3β^35^ (see schematic in Figure 5a). We tested this potential pathway by western blot on mucosal lysates, confirming elevated S9 GSK3β phosphorylation in Sprouty2-null colonic mucosa (Figure 5c). As GSK3β can regulate cytokine levels in some tissues, we asked whether it controls IL-33 expression in the colon and intestine. In both mouse colon epithelial cells (YAMC) and rat intestinal epithelial cells (IEC-6), inhibition of GSK3β using the specific inhibitor CHIR 99021 resulted in increased *IL33* levels (Figure 5d and e). Levels of the epithelial-expressed cytokine, CXCL2, were unaltered demonstrating this response was not a generalized cytokine regulation (Figure 5d and e). Next, since Akt is negatively regulated by Sprouty2^24^ and is responsible for Serine 9 phosphorylation on GSK3β^35^, we also hypothesized that Sprouty2 loss-driven *IL33* induction is through PI3K/Akt. Western blot analysis for Ser473 phospho-Akt identified a significant increase in Spry2^IEKO^ mice (Figure 5b). Furthermore, in YAMC and IEC-6 cells, *IL33* levels were dramatically reduced by PI3K/Akt inhibition (Figure 5d and e). The same findings on IL-33 were observed in primary colon epithelial culture using colonoids (Figure 5f). Furthermore, like IL-33 treatment, GSK3β inhibition was unable to induce tuft and goblet cell markers in colonoids (Figure 5g). Together these data suggest that Sprouty2 repression of PI3K/Akt signaling is permissive for GSK3β activity, and that a baseline inhibition of IL-33 is released when Sprouty2 is repressed by inflammation.

**Figure 5.**
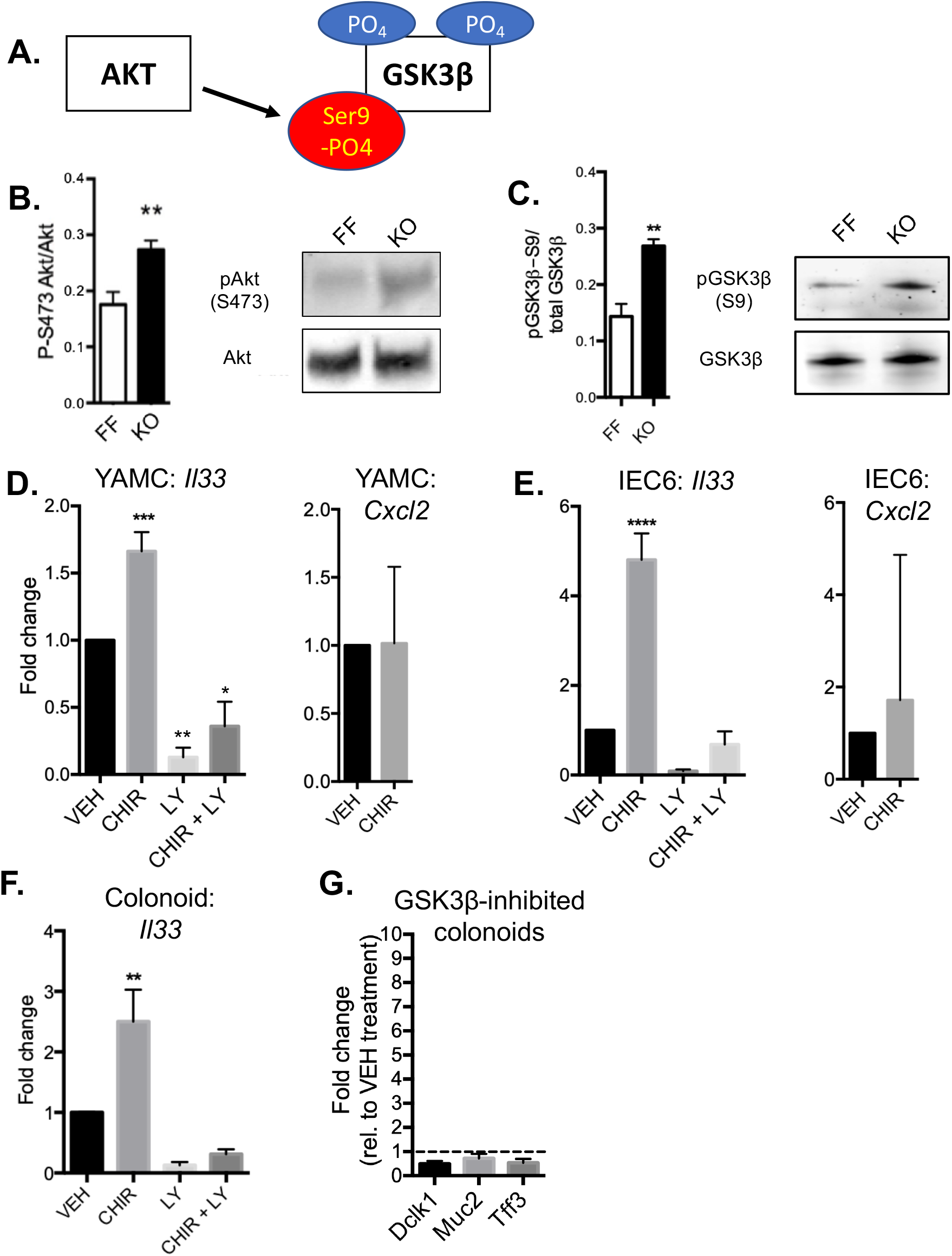
Colonic GSK3beta activity is inhibited by Sprouty2 loss, and promotes IL-33 expression. (A) Schematic of the Akt inhibitory phosphorylation of serine-9 on GSK3β. (B) Serine-473 phospho-Akt and (C) serine-9 phospho-GSK3β levels were measured in colonic homogenates from Spry2^FF^ (FF) and Spry2^IEKO^ mice (KO) by western blot. n=3-5 mice per group. (D and E) The effects of GSK3β inhibition (using CHIR 99021, 3 µM) and/or PI3K/Akt inhibition (using LY294002, 10 µM) for 24 hours on *Il33* and *Cxcl2* expression were measured in mouse colon (YAMC) cells and rat intestinal epithelial (IEC-6) cells. n=5-7 independent experiments per cell line. (F) *Il33* levels were measured in GSK3β or PI3K/Akt inhibited murine colonoids. n=5 independent experiments. (G) *Dclk1*, *Muc2*, and *Tff3* were measured in GSK3β inhibited murine colonoids. n=5 independent experiments. **, p<0.01; ***, *p*<0.001; ****, p<0.0001.

### Spry2^IEKO^ mice are protected against DSS colitis

Goblet and tuft cells play important roles in protecting the colonic epithelium from damage, as demonstrated by spontaneous colitis in *Muc2*^*-/-*^ mice^4,36^ and increased susceptibility to DSS colitis in *Dclk1-*deficient animals^5,28^. Furthermore, IL-33 promotes recovery from DSS colitis^37,38^. This suggests that Spry2^IEKO^ mice, which have increased goblet and tuft cell numbers and elevated IL-33, might be protected from experimental colitis. To test this hypothesis, we subjected Spry2^IEKO^ mice and control littermates to acute DSS (3% in drinking water for 4 d, followed by 3 d chase on standard water). Sprouty2 deletion protected mice from weight loss (Figure 6a), led to improved histology scores (Figure 6b), reduced fecal lipocalin-2 (Figure 6c), reduced colon shortening (Figure 6d), and lowered inflammatory cytokine expression (Figure 6e) following DSS. Apoptosis as measured by TUNEL stain was also significantly reduced in Spry2^IEKO^ colons (Figure 6f). Notably, expression of the goblet cell marker, Muc2, remained elevated following DSS in the Spry2^IEKO^ animals compared to WT (Figure 6h). Furthermore, since Lgr5+ stem cells can be lost in acute inflammatory injury, we measured stem cell markers by qPCR and found that expression of *Lgr5* and *Lrig1* were spared in mice with Sprouty2 deletion in comparison to wild-type (Figure 6g). In contrast, *Bmi1*, which marks reserve stem cells and/or reversion-capable progenitors, was not altered (Figure 6g).

**Figure 6.**
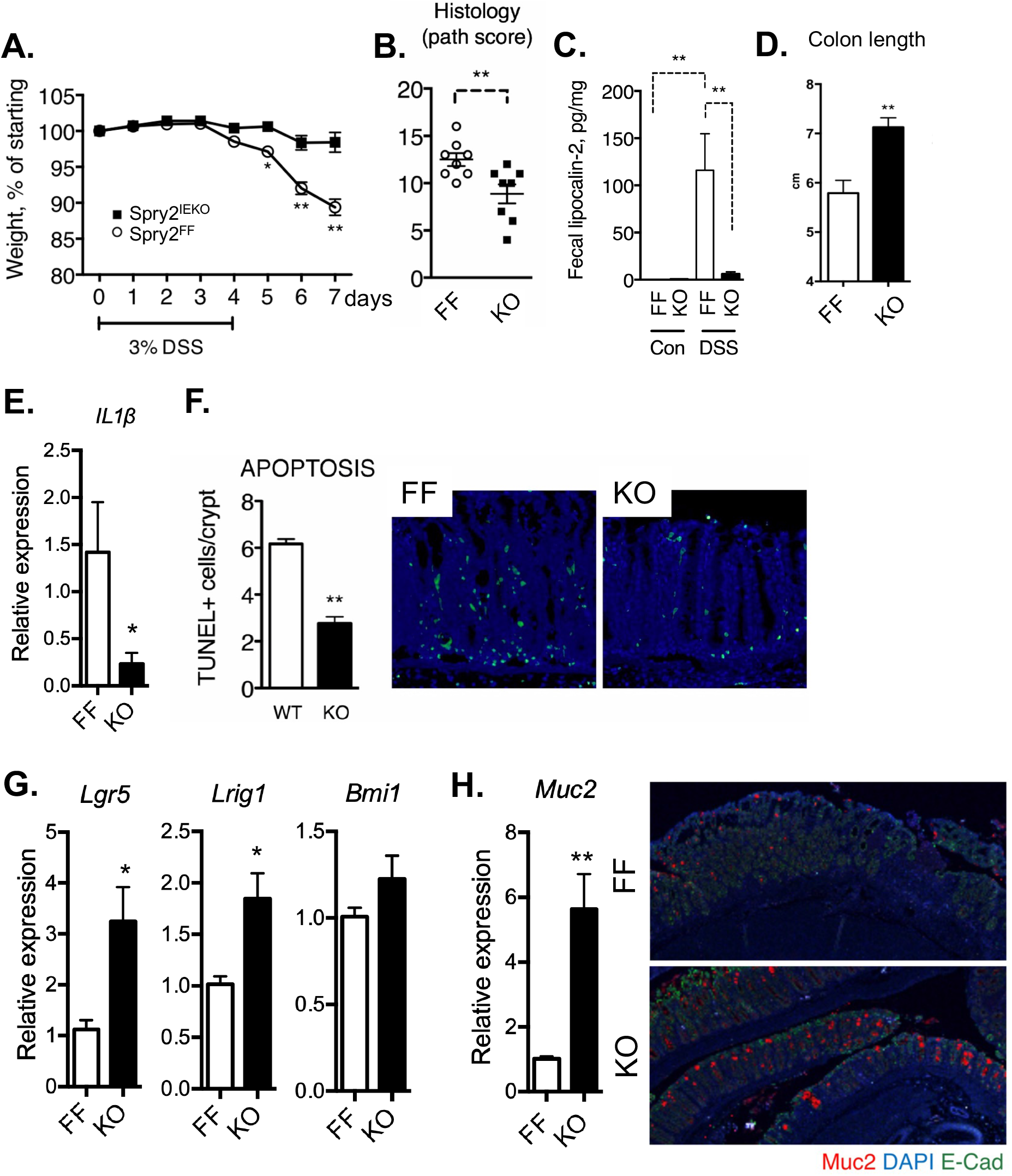
Spry2^IEKO^ mice are protected against DSS colitis. Spry2^FF^ and Spry2^IEKO^ mice were given 3% DSS in drinking water for 4 days, followed by 3 days of plain drinking water to induce colitis; n=11-16 mice per group. (A) Weights were measured daily. (B) Colonic sections were scored by a blinded pathologist. (C) Fecal lipocalin-2, a marker of intestinal inflammation, was measured by ELISA. (D) Colon lengths were measured and (E) expression of *Il1b* was assayed in distal colonic tissue by qPCR between groups in DSS-treated mice at day 7. (F) Apoptosis rate in distal colonic tissue was measured by TUNEL stain. Expression (G) *Lgr5*, *Lrig1*, *Bmi1* and (H) *Muc2* were analyzed by qPCR on collection; colonic sections were subjected to immunofluorescence for Muc2. *, *p*<0.05; **, *p*<0.01.

### Sprouty2 levels are increased in human IBD, and negatively correlate with tuft cell markers

Our results in mice suggest that transient Sprouty2 downregulation by inflammation is a compensatory response to early damage signals (e.g. TNF). To test whether this is consistent in human colon, we generated colonoids from surgical resections of non-IBD patients. Similar to murine colonoids, TNF significantly repressed *SPRY2* expression in these cultures (Figure 7a). To consider whether this mechanism might be defective in IBD, we measured *SPRY2* expression in colon endoscopic mucosal biopsies from pediatric non-IBD and IBD patients with active colonic inflammation. We found elevated expression in both ulcerative colitis and Crohn’s disease patients (Figure 7b). Similar results were obtained from analysis of epithelium from surgical specimens from IBD and control patients (not shown). These data are consistent with findings from Gamo *et al.* demonstrating elevated Sprouty2 expression in adult IBD patients^39^. To ask whether Sprouty2 regulation of secretory cells might be preserved in humans, we correlated *SPRY2* with *DCLK1* and *TRPM5* (tuft cell markers) using GEO dataset GDS3268 consisting of human colonic mucosal biopsies. We found a significant inverse correlation between the tuft markers and Sprouty2 (Figure 7d). Together these data suggest that, while acute inflammation can downregulate Sprouty2 to drive increased numbers of protective secretory cells, in chronic disease this response either does not occur or is lost. Consistent with this, in the IL-10^-/-^chronic colitis model, *Spry2* is elevated in adult animals with active inflammation versus young (pre-colitic) littermates or healthy wild-type controls (Figure 7e). A failure to down-regulate Sprouty2 in response to early damage signals may be a factor in IBD development (Figure 8).

**Figure 7.**
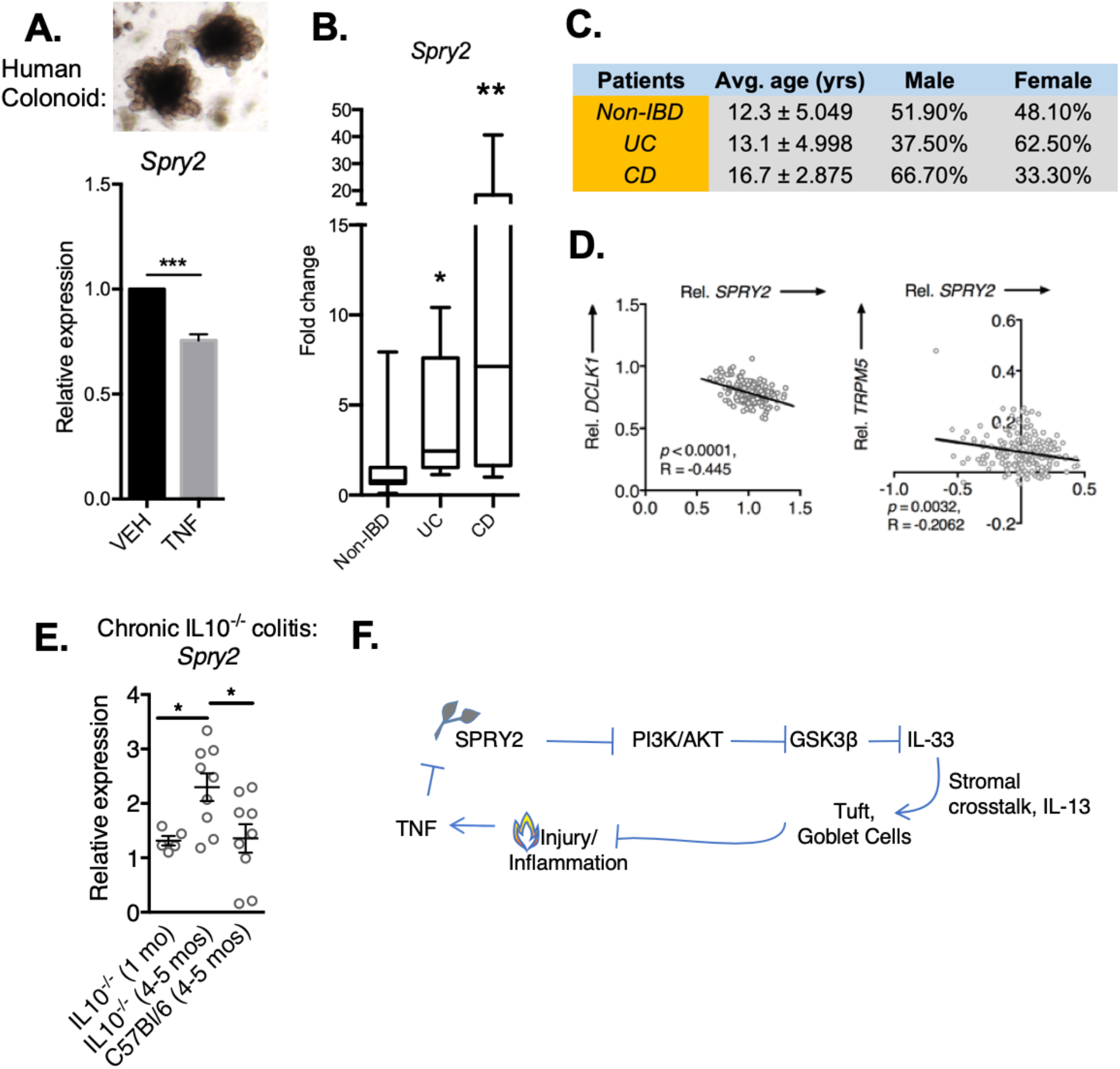
Sprouty2 is elevated in human IBD, and expression levels negatively correlate with tuft cell markers. Human colonoids generated from surgical specimens were treated with TNF (10 ng/ml) for 24 hours and *SPRY2* expression was analyzed by qPCR. n=4 independent experiments per group. Samples from patient mucosal biopsies from non-IBD (n=27), or Crohn’s disease (n=6), and ulcerative colitis patients (n=8) with active inflammation were analyzed for *SPRY2* expression; patient characteristics shown in (C). (D) The publicly available GEO dataset (GDS3268) containing RNA sequencing data from colon biopsies was analyzed for *SPRY2* and tuft cell marker (*DCLK1, TRPM5*) expression, and correlation analysis was performed revealing a significant negative correlation between Sprouty2 and tuft cell markers. (E) Distal colons from IL10^-/-^mice (n=5-9 per group) before onset of colitis (1 month of age) and post-onset of chronic colitis (4-5 months of age) were assayed by qPCR for *Spry2* expression, compred to healthy wild-type C57Bl/6 animals. *, p <0.05; **, p <0.01; ***, p <0.001. (F) Proposed model of Sprouty2 regulation of epithelial IL-33 and secretory cell development in response to colonic inflammation: Acute injury/inflammation drive TNF expression, thus repressing Sprouty2 expression in the epithelium. This removes basal inhibition on PI3K/Akt, leading to Akt-mediated inhibition of GSK3β, promoting epithelial IL-33 expression. Epithelial-produced IL-33 induces IL-13 in the stroma, which in turn signals back to the epithelium to promote tuft and goblet cell expansion as a compensatory mechanism to limit acute inflammation and promote injury resolution. Loss of this dynamic differentiation mechanism might contribute to chronic disease.

## Discussion

Maintenance of homeostasis and effective responses to injury in the colonic epithelium require a well-regulated balance of different epithelial cell types including stem, absorptive, and secretory cells. Epithelial-, stromal-, and immune-derived factors direct the development of these cell types^40–44^. However, the precise mechanisms that integrate these signals in the local environment to alter the cellular composition and function of the epithelium are incompletely understood.

In this study, we sought to define the role of Sprouty2 in controlling these processes. Growth factor receptor-initiated signaling, including PI3K and MAPK cascades, is indispensable for intestinal homeostasis^45^ and directs intestinal epithelial cell development and differentiation^9,11,46^. Sprouty2 can negatively regulate PI3K and MAPK cascades, and is itself regulated by growth factor signals^22,26^. Here we show that it is also repressed by inflammation both in vitro and in vivo (Figures 1 and 7), and thus may play a role in regulating the colonic response to inflammation and injury.

The use of intestinal epithelial-specific Sprouty2 deleted mice (Spry2^IEKO^) allowed us to isolate the effects of Sprouty2 loss from the broader inflammatory milieu of colitis, allowing us to directly test its function. Spry2^IEKO^ mice showed increased numbers of secretory tuft and goblet cells, but not enteroendocrine cells or absorptive enterocytes. We speculate this induction may be a compensatory skewing towards cell types that are protective in mouse models of colitis^4,5,47^ and in patients with IBD^48^. Under this scenario, Sprouty2 is constitutively expressed in homeostasis, but is repressed by a danger signal (e.g. TNF) to enable a rapid response to insult.

To identify downstream mediators driving these effects, we screened for factors known to skew cells towards secretory lineages. This led to the novel finding that Sprouty2 deletion induces IL-33 production in colonic epithelial cells. IL-33 is expressed by epithelial, stromal, and immune cells, though at homeostasis, epithelial expression is normally low^49,50^. IL-33 regulation has predominantly been studied in immune cells^51^ and control of its expression in the intestinal epithelial compartment is not well understood. It has documented roles as a nuclear alarmin and an intracellular regulator of gene function, as well as potentially serving as a traditional secreted cytokine^52^. We found that Sprouty2-regulated IL-33 (Figure 3) initiated an epithelial-stromal circuit (Figure 4), suggesting that it is the secreted IL-33 form functioning here.

The role of IL-33 in colonic disease is controversial, as some studies suggest it is pathogenic^53,54^ while others have found it is associated with improved outcomes in colitis^38^. These different results likely depend on the cellular source of IL-33 and timing of when the cytokine is expressed in the course of disease^55^. Our data suggest that when IL-33 is expressed in the epithelium prior to the onset of disease, it conditions the epithelium against future insult (such as the chemical damage in DSS colitis). Using colonoids from Spry2^IEKO^ and littermate controls, we show that epithelial loss of Sprouty2 directly regulates epithelial IL-33 expression and its impact on secretory cell differentiation is due to extra-epithelial interactions of the cytokine.

While elevated epithelial IL-33 was maintained in Spry2^IEKO^ colonoids following passage, tuft and goblet cell lineage markers were not. Furthermore, exogenous IL-33 did not induce tuft or goblet cell markers. This suggests IL-33 does not promote secretory differentiation by direct epithelial-intrinsic signaling, but rather requires interaction with stromal or recruited cells.

Recent work has shown that IL-33 can activate innate lymphoid cells to produce IL-13^52,56^. Waddell *et al.* demonstrated that this cytokine then reciprocally acts on the intestinal epithelium to promote goblet cell expansion^34^. Consistent with this, we found elevated numbers of IL-13^+^ cells in Spry2^IEKO^ colons; furthermore, IL-13 but not IL-33 induced tuft and goblet cell markers in wild-type colonoids. Importantly, we also confirmed a requirement for IL-13 by treating IL-13KO and WT mice with IL-33, showing induction of tuft and goblet markers only in WT mice.

An important advance from this work is the identification of Sprouty2-regulated intracellular signaling pathways in the colonic epithelium. Spry2^IEKO^ mice had elevated levels of Akt activation and phosphorylation of the inhibitory Serine 9 on GSK3β. Akt mediates this phosphorylation site^35^. This suggests a model where TNF-induced loss of Sprouty2 releases baseline inhibition on Akt signaling, allowing for Serine 9 phosphorylation on GSK3β and induction of IL-33. Our results might indicate a previously unrecognized role for Wnt signaling in the control of intestinal IL-33, though it should be noted that GSK3β could work through other targets and thus a Wnt connection will have to be formally tested in future studies. GSK3β and Wnt signaling are known to influence the balance of secretory cells in the intestine^57^, and our work reveals Sprouty2 as a novel upstream regulator controlled by inflammation and links this pathway to induction of IL-33 (Figure 5). Previous reports have found that IL-33 regulates Akt signaling^58^. However, the impact of Akt in control of IL-33 signaling in intestinal tissues has not previously been tested. In accord with Ivanov *et al.* showing that Akt positively regulates IL-33 signaling in human fibroblasts^59^, we found that inhibition of the PI3K/Akt cascade repressed basal IL-33 levels from the colonic epithelium. We cannot formally exclude the possibility that altered GSK3β signaling in Spry2^IEKO^ mice also provides feedback to Akt to heighten IL-33 production, but at minimum our in vitro studies do show that IL-33 is regulated by PI3K.

The epithelial switches controlling IL-33 production in the colonic epithelium, especially in the context of inflammation, are not well understood. Here we have identified Sprouty2 as a crucial integrator of extracellular stimuli (i.e. TNF) that modulates downstream signaling pathways controlling epithelial IL-33 and thus tuft and goblet cell specification. Tuft cells regulate an inflammatory circuit and both tuft and goblet cells protect against colitis in mice^4,5,28^.

Furthermore, when these cell types are genetically targeted in mice, experimental colitis is exacerbated. Together these observations suggest the possibility that the Sprouty2 circuit described here might be compromised in IBD, a possibility supported by our data showing a significant increase in *SPRY2* in colonic specimens from IBD patients. As patients with elevated type 2 immune response signatures (such as IL-13) have better outcomes in IBD^48^, an important next step will be to understand if Sprouty2 expression or its regulation by TNF is a valid biomarker for IBD severity.

Elevated Sprouty2 in IBD was an unexpected finding as mucosal biopsies from ulcerative colitis patients (and to a lesser extent Crohn’s disease) can in some studies express elevated IL-33^60^. However most biopsy samples contain epithelial, mesenchymal, and immune cells making determination of the IL-33 source in these studies unclear, whereas Sprouty2 is primarily epithelial. The source, processing (mature vs full-length forms), and localization of IL-33 within the colon likely play a key role in determining the outcome of its effects on tuft and goblet cell development; and we have found that epithelial IL-33 is specifically protective.

In summary, we have demonstrated that Sprouty2 is an inflammation-responsive protein in the colonic epithelium that maintains a cytokine circuit to control tuft and goblet cell expression. This represents a novel way in which the epithelium regulates Akt and GSK3β signaling to produce IL-33, and thus alter the composition of the epithelial barrier. This negative feedback mechanism appears to be protective against colonic inflammation.

## Materials & Methods

### Animal experiments

C57Bl/6 mice obtained from Jackson Laboratory aged 8-10 weeks were used for experiments. Mice with an intestinal epithelial-specific deletion of Sprouty2 (VillinCre;Spry2^flox/flox^) were generated by crossing VillinCre animals with mice harboring loxP-flanked Spry2 (Spry2^flox/flox^). VillinCre;Spry2^flox/flox^ and Spry2^flox/flox^ littermate controls aged 8-10 weeks were used for experiments. IL-13^-/-^mice and BALB/c controls aged 6-12 weeks were given daily i.p. injections of 0.4 µg rIL-33 for 4 days as previously described^34^. For acute colitis, mice were given 3% (w/v) dextran sodium sulfate (DSS) in drinking water for 4 days, followed by 3 days without drinking water.

### Real-time PCR

RNA from cells and tissue was collected using on-column RNA isolation and purification (OMEGA Biotek), and cDNA generated with a high-capacity cDNA reverse transcriptase kit (Applied Biosystems, 4368814). Quantitative analysis of expression was performed using TaqMan assays (Spry2 (Mm00442344_m1, Hs01921749_s1), Dclk1 (Mm00444950_m1), Trpm5 (Mm01129032_m1), Il25 (Mm00499822_m1), IL33 (Mm00505403_m1, Rn01759835_m1), Il10 (Mm01288386_m1), Il4 (Mm00445259_m1), ChgA (Mm00514341_m1), Muc2 (Mm01276696_m1), Tff3 (Mm00495590_m1), Lgr5 (Mm00438890_m1), Lrig1 (Mm00456116_m1), Bmi1 (Mm03053308_g1), Car2 (Mm00501576_m1), Aqp8 (Mm00431846_m1), Alox5 (Mm01182747_m1), Pou2f3 (Mm00478293_m1), Il1β (Mm00434228_m1), Tnf (Mm00443258_m1), Cxcl2 (Mm00436450_m1, Rn00586403_m1), and Hprt (Mm03024075_m1, Rn01527840_m1, Hs02800695_m1)) on an Applied Biosystems StepOne thermocycler. Fold change was calculated using the 2^-ΔΔCt^ method. Results are expressed as average fold change in gene expression relative to control or non-treatment group using *Hprt* as the reference gene.

### RNAScope In-situ hybridization

Distal colonic section were probed using RNAScope probes Mm-Spry2 (Advanced Cell Diagnostics, #425061) or Mm-Il13 (Advanced Cell Diagnostics, #312291) using the RNAScope 2.5 HD Detection system (Advanced Cell Diagnostics, #322310) according to manufacturer-provided protocol.

### Western blotting and proteome profiler

Protein lysates from cells and tissue were collected and lysed in RIPA buffer with Halt Protease inhibitor cocktail (Thermo Scientific, #1861278), and phosphatase inhibitor cocktails 2 and 3 (Sigma, P5726 and P0044)^61^. Protein concentration was determined by DC protein assay (Bio-Rad, #500). For proteome profiling, samples were analyzed according to manufacturer instructions using the Phospho-MAPK Array (R&D Systems, #893909). For western blots, 30 µg protein/condition were separated by SDS-PAGE (Thermo Scientific, NW0412A) and transferred to nitrocellulose membrane. Membranes were blocked with 5% milk and probed with 1:1000 Sprouty2 (Sigma, #AV50523) overnight at 4°C, 1:1000 total and phospho-GSK3beta (S9) (Cell Signaling, #12456, #5558, respectively), 1:1000 total and phospho-Akt (S473) (Cell Signaling, #2920, #4060, respectively) or 1:10,000 mouse anti-Actin (Sigma, A1978) for 1 hour at room temperature, followed by 1:20,000 IRDye-conjugated donkey anti-rabbit (LI-COR, #926-68023) and donkey anti-mouse (LI-COR, #926-32212) for 1 hour at room temperature and quantification on an Odyssey imager (LI-COR).

### ELISA

Fecal lipocalin-2 levels in distal colon fecal contents were analyzed using Mouse Lipocalin-2 DuoSet ELISA (R&D Systems, DY1857) according to manufacturer provided instructions. IL-33 in colonic mucosa of distal colon was analyzed using Mouse IL-33 DuoSet ELISA (R&D Systems, DY3626) according to manufacturer instructions.

### Immunofluorescence staining

Distal colon sections (5 µm), from tissue fixed with 4% formaldehyde overnight and paraffin embedded, were dewaxed and blocked with 10% goat serum for 1 hour at room temperature. This was followed by incubation with primary antibody against DCLK1 (Abgent, #AP7219B), ChgA (ImmunoStar, #20085), Muc2 (Abcam, Ab76774), or E-cadherin (BDBiosciences, #610181) overnight at 4 °C. Cells were washed and incubated with secondary Alexa Fluor-488 (Life Technologies) or Alexa Fluor-555 (Life Technologies) for 1 hour at room temperature followed by mounting with Vectashield mounting media including DAPI (Vector Labs, H-1500).

### Barrier function

To test intestinal permeability, mice were gavaged with 200 µl of 22 mg/ml 4 kDa-FITC dextran (Sigma, #46944) in normal saline 4 hours before sacrifice. Blood was collected by cardiac puncture into EDTA containing tubes, and placed on ice. Analysis for FITC fluorescence was performed on a Molecular Devices SpectraMax M3 fluorescence plate reader.

### Cell lines and colonoids

For in vitro experiments, immortalized young adult mouse colon (YAMC) cells and immortalized rat small intestinal cells (IEC-6) were grown to 90% confluency before use in experiments. Colonoids were generated and passaged from mouse colon and human colon using previously established protocols^9,62,63^. Briefly, crypts were isolated and embedded into Matrigel (BD Biosciences), then overlaid with mouse or human growth media. Cultures were passaged every 7-10 days. Cultures were treated with the indicated concentrations of murine TNF (Peprotech, #315-01), human TNF (Peprotech, #300-01), mouse IL-33 (Peprotech, #210-33), or mouse IL-13 (Peprotech, #210-13) for the time specified before collection. TUNEL stain was performed as previously described^64^ on distal colonic sections from DSS-treated mice.

### Human tissue

Colonic tissue from endoscopic biopsies or gross surgical specimens were obtained from pediatric patients either with or without IBD at Children’s Hospital Los Angeles. Samples were placed immediately in RNALater (Thermo Fisher Scientific, AM7021) or used for colonoid generation.

### Statistics

Statistical analyses and plots were generated using Prism 6 (GraphPad Software). Mean +/−SEM is depicted in dot and bar graphs. Student’s t test or ANOVA with Tukey post-hoc test to correct for multiple comparisons were used to determine statistical differences, as appropriate. Statistical significance was assigned to *p* < 0.05, and indicated in figure legends.

### Study Approval

All animal use was approved and monitored by the Children’s Hospital Los Angeles Institutional Animal Care and Use Committee (Animal Welfare Assurance #A3276-01) or the Cincinnati Children’s Hospital Medical Center Institutional Animal Care and Use Committee (Animal Welfare Assurance #A3108-01). Mice were housed under standard conditions with ad libitum water and chow access in the AAALAC-accredited animal care facilities at Children’s Hospital Los Angeles or Cincinnati Children’s Hospital Medical Center. Human tissue was collected after written informed consent was obtained, under approved Institutional Review Board CCI-13-00287 and CCI-09-00093, respectively, at Children’s Hospital Los Angeles.

## Author Contributions

MAS: conceived and designed experiments, performed experiments, analysis/interpretation of data, wrote the manuscript. JJH, CYU, KLA, DA, KK, JKB, EBB: performed experiments, analyzed data, revised manuscript. AW, SG, KMM: performed experiments, revised manuscript. MKW: analyzed data, revised manuscript. TCG, DW: conceived experiments, revised manuscript. MJR: conceived and designed experiments, revised manuscript. MRF: study supervision, obtained funding, performed experiments, conceived and designed experiments, analysis/interpretation of data, wrote manuscript.

## Acknowledgements

This work was supported by NIH grants R01DK095004 and R01DK119694 (MRF) and R01DK117119 (MJR), a Research Fellowship Award (AW) and a Career Development Award from the Crohn’s and Colitis Foundation (MAS), and funding from CURE for IBD.

